# Genome sequences of *Arthrobacter* spp. that use a modified sulfoglycolytic Embden-Meyerhof-Parnas pathway

**DOI:** 10.1101/2021.11.24.469816

**Authors:** Arashdeep Kaur, Phillip L. van der Peet, Janice W.-Y. Mui, Marion Herisse, Sacha Pidot, Spencer J. Williams

## Abstract

**Background:** Sulfoglycolysis pathways enable the breakdown of the sulfosugar sulfoquinovose and environmental recycling of its carbon and sulfur content. Several pathways exist for the breakdown of sulfoquinovose that usually lead to production of C3-sulfonates (sulfolactate and 2,3-dihydroxypropanesulfonate) that are excreted and sustain secondary bacterial communities. The prototypical sulfoglycolytic pathway is a variant of the classical Embden-Meyerhof-Parnas pathway that has been described in gram-negative *Escherichia coli* and results in production of 2,3-dihydroxypropanesulfonate.

**Results:** We used enrichment cultures to discover new sulfoglycolytic bacteria from Australian soil samples. Two gram-positive *Arthrobacter* spp. were isolated that produced sulfolactate as the metabolic end-product. Genome sequences identified a modified sulfoglycolytic Embden-Meyerhof-Parnas (sulfo-EMP) gene cluster that retained the core sulfoglycolysis genes encoding metabolic enzymes, but featuring the replacement of the gene encoding sulfolactaldehyde (SLA) reductase with SLA dehydrogenase, and the absence of sulfoquinovosidase and sulfoquinovose mutarotase genes. The gene clusters were broadly conserved across a range of other sequenced Actinobacteria.

**Conclusions:** We report the first gram-positive soil bacteria that utilize sulfo-EMP pathways to metabolize SQ. Excretion of sulfolactate is consistent with an aerobic saprophytic lifestyle. This work broadens our knowledge of the sulfo-EMP pathway to include soil bacteria.

## Background

Sulfoquinovose (SQ; 6-deoxy-6-sulfo-D-glucose) is a sulfosugar produced by photosynthetic organisms [1]. It is primarily found as the headgroup of the sulfoglycolipid sulfoquinovosyl diacylglycerol (SQDG) in photosynthetic tissues and membranes in plants, algae and cyanobacteria [1, 2]. The annual global production of SQ is estimated at 10^10^ tons per annum [3], and thus the degradation of this sulfosugar is an important arm of the global biogeochemical cycle. The degradation of sulfoquinovose occurs through pathways of sulfoglycolysis and provides access to its constituent carbon and generates ATP and reducing equivalents (NADH/NADPH) [4]. The sulfoglycolytic Embden-Meyerhof-Parnas (sulfo-EMP) [5, 6], Entner-Doudoroff (sulfo-ED) [7] and sulfofructose transaldolase (sulfo-SFT) [8, 9] pathways cleave the 6-carbon chain of SQ into two 3-carbon fragments, one of which is utilized in primary metabolism, while the other containing the sulfonate group is converted to either sulfolactate (SL) or 2,3-dihydroxypropanesulfonate (DHPS) and excreted. The sulfoglycolytic sulfoquinovose monooxygenase (sulfo-SMO) pathway results in cleavage of the C-S bond of sulfoquinovose and leads to production of glucose, and thus enables the complete breakdown of the SQ molecule [10]. The genes encoding these sulfoglycolytic pathways are found within clusters that typically contain genes encoding proteins for the import of SQ or its glycosides, the export of the end-products, SL or DHPS, and a specialized glycoside hydrolase termed a sulfoquinovosidase (SQase) [11, 12] that can cleave SQ glycosides to release SQ that can undergo sulfoglycolysis.

We report here the use of sequential enrichment culturing using minimal media containing SQ as sole carbon source to isolate new sulfoglycolytic bacteria from soil. We identify two *Arthrobacter* sp. strains, AK01 and AK04, and demonstrate that they grow on SQ and secrete SL into the growth media. We present the draft genome sequences of AK01 and AK04 that reveals that these *Arthrobacter* sp. contain a gene cluster encoding a sulfoglycolytic Embden-Meyerhof-Parnas pathway that differs from the prototypical pathway of *E. coli* through the lack of an identifiable candidate SQase, the replacement of SLA reductase with SLA dehydrogenase, and the presence of ABC transporters and TauE permeases, variations that are present within other sequenced Actinobacteria.

## Results and Discussion

### Isolation and characterization of sulfoglycolytic bacteria

Bacteria able to grow on SQ as sole carbon source were selected by using soil samples (collected from the University of Melbourne, Parkville campus) to inoculate a vitamin-enriched minimal media containing SQ as sole carbon source. Sequential subculturing into fresh SQ-minimal media, followed by plating onto LB agar and then regrowth in SQ-minimal media led to isolation of strains AK01 and AK04 that possessed a short rod-like appearance (**Fig. 1**). Strains AK01 and AK04 grew robustly on SQ with peak growth rates of 0.129 and 0.081 A_600_/min and achieved stationary phase after approximately 25 and 40 h, respectively (**Fig. 2a**,**c**). The absorbance at stationary phase for cultures grown on SQ were approximately half of that for cultures grown on equimolar glucose. ^13^C-NMR analysis of culture medium of AK01 and AK04 grown on ^13^C_6_-SQ (7.7 mM) media gave three signals corresponding to ^13^C_3_-SL (**Fig. 2b**,**d, Table 1**). ^13^C NMR analysis of spent culture confirmed that substrate SQ is completely consumed by bacteria and metabolised to SL.

**Figure 1.**
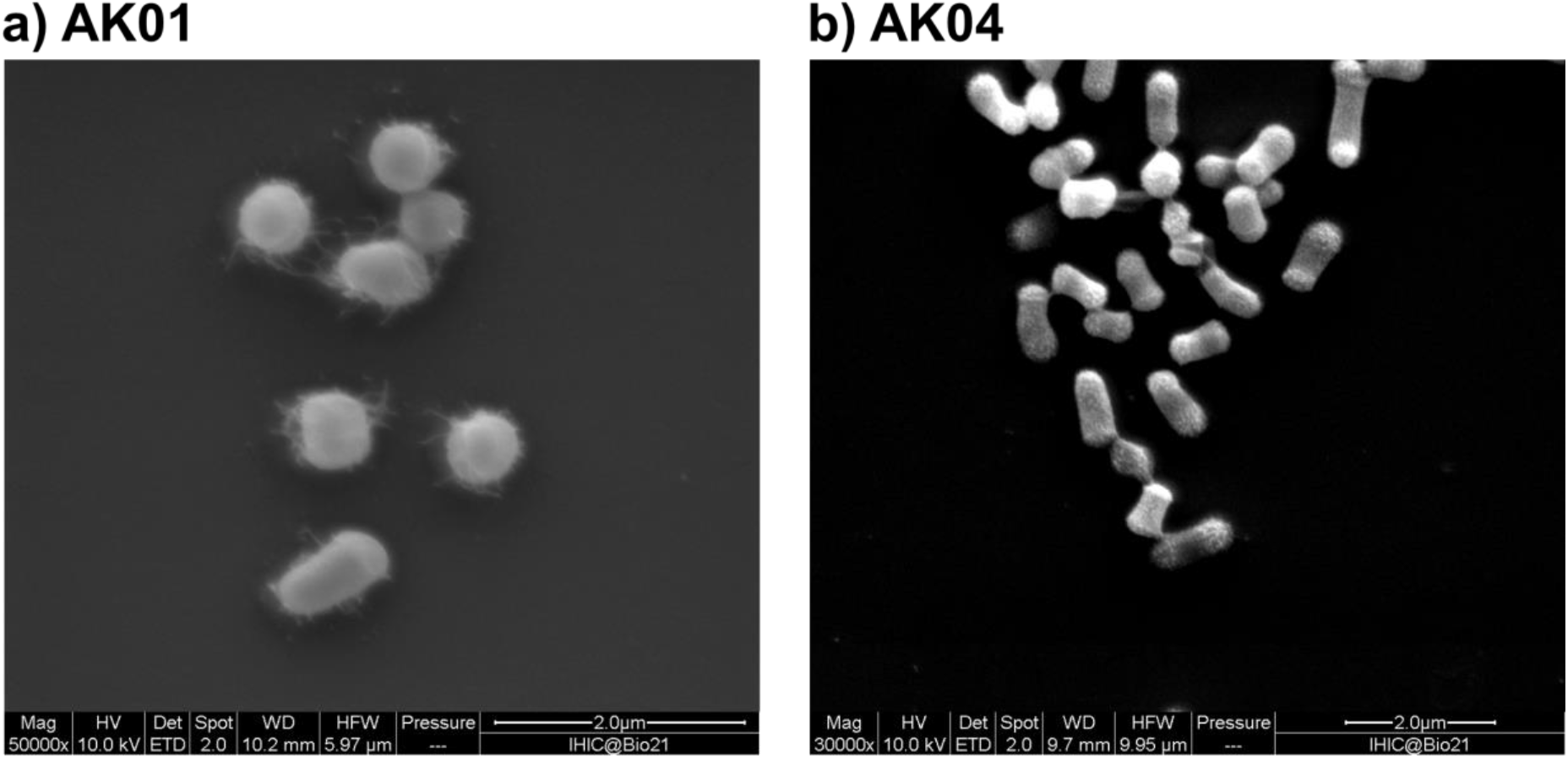
Scanning electron microscopy image of *Arthrobacter* sp. a) strain AK01, b) strain AK04. Cell morphology was examined using a scanning electron microscope (Quanta 200 ESEM). Cells were grown in LB media for 3 days, fixed in 0.05% glutaraldehyde in 0.1 M phosphate buffer (pH 7.4), then in 2.5% glutaraldehyde in 0.1 M phosphate buffer (pH 7.4) and allowed to react for 20 min. Fixed cells were adhered onto poly-lysine coated slides and rinsed with water 3 times, then dehydrated by soaking in an ascending ethanol gradient (20-100%). The sample was critical point dried using a Leica CPD3000 and gold coated to thickness of 5 nm using Safematic CCU-010 compact coating unit. Images are at approximately 50,000x magnification with scale bar shown.

**Figure 2.**
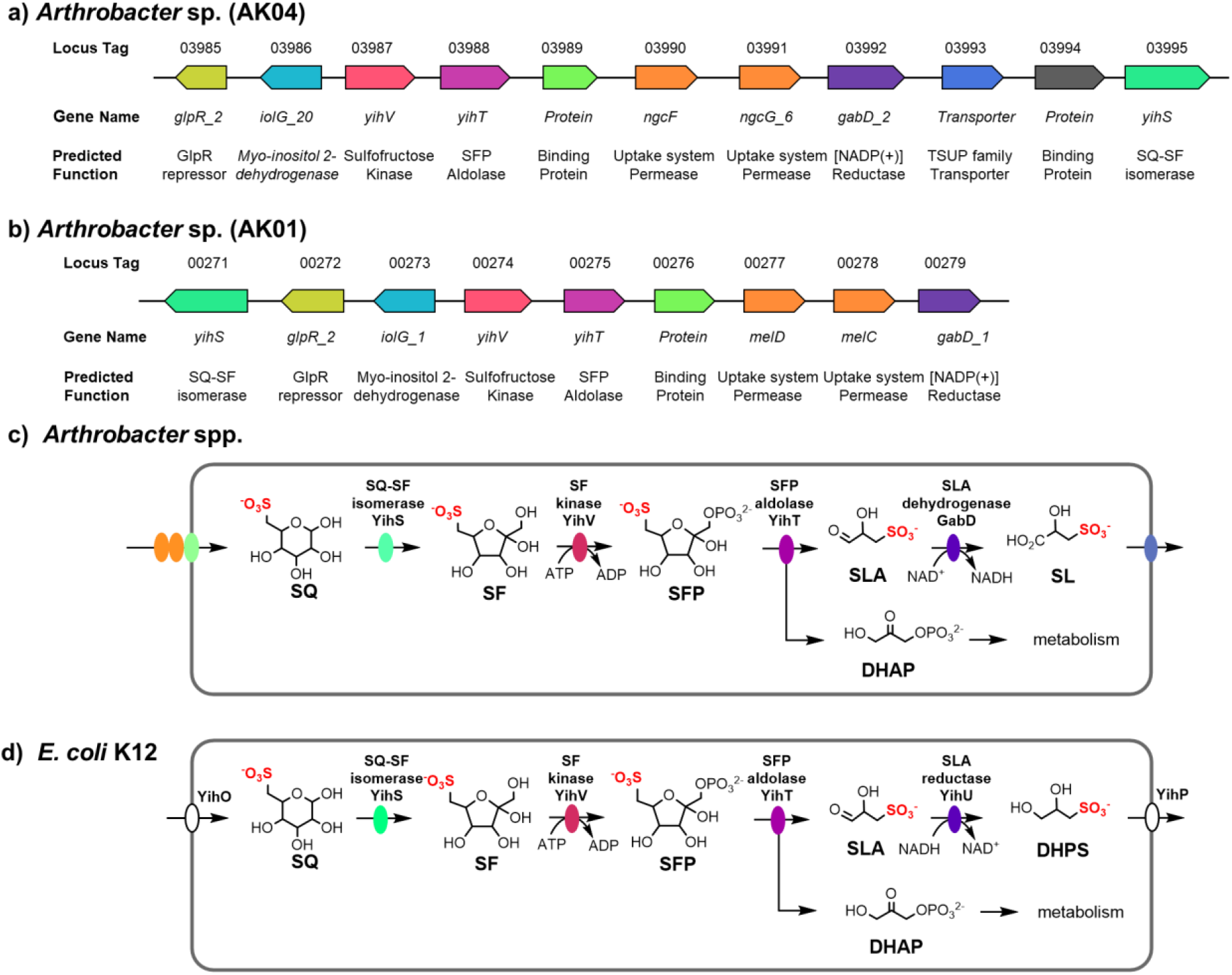
Proposed sulfoglycolytic Embden-Meyerhof-Parnas (sulfo-EMP) pathway for *Arthrobacter* spp. (a) Gene cluster encoding the sulfo-EMP pathway for *Arthrobacter* sp. AK04 (b) Gene cluster encoding the sulfo-EMP pathway for *Arthrobacter* sp. AK01 (c) Proposed sulfo-EMP pathway for *Arthrobacter* spp. (d) Comparison with EMP pathway for *E. coli* K12.

**Table 1.** ^13^C NMR (125 MHz) data of ^13^C_3_-SL produced as metabolite from ^13^C_6_-SQ. Samples contain 50 % D_2_O to allow frequency lock.

| chemical shift<br>(d, ppm) | multiplicity | coupling constant<br>(Hz) | assignment |
| --- | --- | --- | --- |
| 55.19 | d | $^1J_{\text{C1,C2}} = 36.9$ | C3 |
| 62.42 | s | NA | --- |
| 68.85 | dd | $^1J_{\text{C2,C3}} = 52.7$<br>$^1J_{\text{C1,C2}} = 38.0$ | C2 |
| 72.01 | s | NA | --- |
| 178.69 | d | $^1J_{\text{C2,C3}} = 54.6$ | C1 |

To investigate the genetic basis for SQ consumption by AK01 and AK04, DNA extracted from these bacteria were sequenced using the Illumina NextSeq platform. **Table 2** shows the key features of the two draft genomes. On the basis of 16S rRNA gene sequence analysis the organisms were assigned as *Arthrobacter* sp. The 16S rRNA genes of the two strains were 98.8% identical over 1520 bp, which suggests that these organisms are of the same species [13]. However, average nucleotide identity (ANI) between the two organisms is only 76.5%, conflicting with the 16S rDNA gene results and suggesting that these two organisms differ sufficiently to be considered separate species [14].

**Table 2.** Key features of the AK01 (GenBank accession: SAMN23041292) and AK04 (GenBank accession: AK01, SAMN23041293) draft genome assemblies.

|  | **AK01** | **AK04** |
| --- | --- | --- |
| Genome size | 5,105,913 | 4,700,363 |
| Number of contigs | 169 | 130 |
| GC % | 62.87 | 65.51 |
| Number of ORFs | 4734 | 4318 |
| Number of putative genes | 4669 | 4261 |
| Number of putative tRNA | 59 | 53 |
| Number of putative rRNA | 5 | 3 |
| Number of putative tmRNA | 1 | 1 |
| Number of genes assigned a function (%) | 2478 (53%) | 2219 (52%) |

### Genomic features related to SQ metabolism

Genome analysis of *Arthrobacter* spp. AK01 and AK04 revealed a cluster of genes that were assigned as encoding SQ degradation through the sulfo-EMP pathway (**Fig. 3a**,**b**). The prototypical sulfo-EMP pathway was identified in *E. coli* and involves a 10-gene cluster (*yihOPQRSTUV, csqR*) [5]. In *E. coli* these genes encode a transcription factor (CsqR) [15], putative transmembrane proteins for the import of SQ and export of the end-product of the pathway, DHPS (YihO, YihP). The enzymatic steps involve a sulfoquinovosidase (YihQ) for cleavage of SQ glycosides [12], sulfoquinovose mutarotase (YihR) for interconversion of SQ anomers [16], SQ-sulfofructose (SF) isomerase (YihS) that interconverts SQ, SF and sulforhamnose,[6] an ATP-dependent sulfofructose kinase (YihV) that converts SF to SF-1-phosphate (SFP) [6], sulfofructose-1-phosphate aldolase (YihT) which converts SFP to SLA and dihydroxyacetone phosphate [6], and a NADH-dependent sulfolactaldehyde reductase (YihU) to convert SLA to DHPS [17], which is excreted into the growth media (**Fig. 3d**).

**Figure 3.**
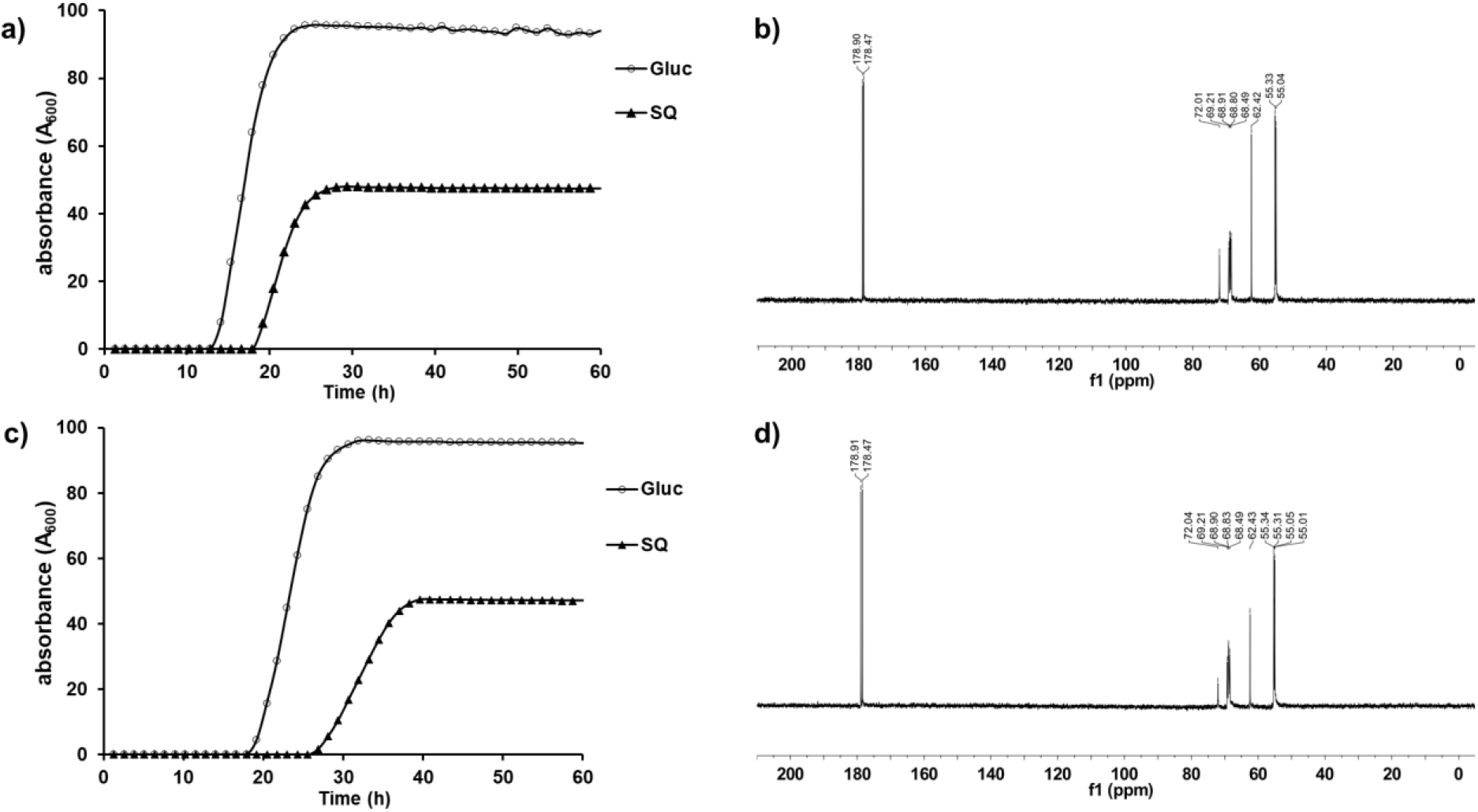
Growth curves of *Arthrobacter* strains a) AK01 and c) AK04 grown on minimal salts media containing 5 mM glucose or SQ. ^13^C NMR (500 MHz) spectra of spent culture media of *Arthrobacter* strains b) AK01 and d) AK04 grown on ^13^C_6_-SQ (7.7 mM).

Both *Arthrobacter* sp. contained genes encoding SQ-SF isomerase (YihS), SF kinase (YihV), and SFP aldolase (YihT). Consistent with the excretion of SL into the growth media, both strains lacked an SLA reductase homologue, but instead contained an SLA dehydrogenase homologue, annotated as GabD. GabD homologues within sulfoglycolytic gene clusters have been identified for bacteria that utilize the sulfo-ED [7, 18] and sulfo-SFT [8, 9] pathways, and in *Pseudomonas putida* SQ1 there is an NAD^+^/NADP^+^-dependent SLA dehydrogenase [7]. A proposed pathway for SQ metabolism in these *Arthrobacter* spp. is shown in **Figure 3c**. AK01 and AK04 represent the first characterized examples of sulfoglycolytic bacteria that use a sulfo-EMP pathway with a GabD SLA dehydrogenase. *E. coli* and related Enterobacteriacae that contain SLA reductases are facultative anaerobes, and the presence of a reducing pathway for excretion of DHPS may support their anaerobic lifestyle. On the other hand, *Arthrobacter* are normally considered aerobes [19] (although anaerobic *Arthrobacter* have been described [20]), which is consistent with an oxidative pathway for excretion of SL.

SQ import and DHPS/SL export is poorly characterized, but several different strategies have been identified across sulfoglycolytic pathways. Both AK01 and AK04 contain genes annotated as ABC transporter cassettes and solute binding proteins. ABC transporter systems have been identified in *Agrobacterium tumefaciens* C58 (which uses the sulfo-SMO pathway) {Sharma, 2021 #8628} and *R. leguminosarum* SRDI858 (which utilizes a sulfo-ED pathway) [18]. The *A. tumefaciens* solute binding protein binds SQGro with high affinity [10]. We therefore propose that these *Arthrobacter* isolates utilize the solute binding protein and an ABC transporter system to import SQ or its glycosides. Strain AK04 also contains a TauE permease (a member of the 4-toluene sulfonate uptake permease (TSUP) system) [21]. TSUP proteins are poorly characterized permeases that are suggested to be involved with the transport of sulfur-containing organic compounds. TauE of *Cupriavidus necator* H16 is proposed to be involved in the export of sulfolactate [22], a function that is consistent with the excretion of SL by AK04. The absence of an obvious permease candidate in the AK01 gene cluster suggests that another protein may adopt this function in this strain.

Other key differences with *E. coli* includes the lack of identifiable SQ mutarotase and SQase encoding genes. Interestingly, the isolated strains could grow on the simple SQ glycoside, methyl α-sulfoquinovoside. This suggests that they may harbor an unidentified SQase without homology to known SQases.

A search for organisms with gene clusters related to strains AK01 and AK04 led to identification of other *Arthrobacter* strains with syntenic or closely related sulfo-EMP gene clusters (**Fig. 4**). Other *Arthrobacter* spp. were identified that contained gene clusters with architectures identical to AK01; and both *Arthrobacter* spp. and *Pseudarthrobacter* spp. were identified with gene clusters identical to AK04. Non-identical but closely syntenic sulfo-EMP gene clusters were observed in select Actinobacteria including other *Arthrobacter* spp. and *Microbacterium* sp. No. 7 (both order Micrococcales), and a more distantly related cluster in the actinobacteria *Acrocarpospora corrugata* (order Streptosporangiales) and *Phytohabitans houttuyneae* (order Micromonosporales).

**Figure 4.**
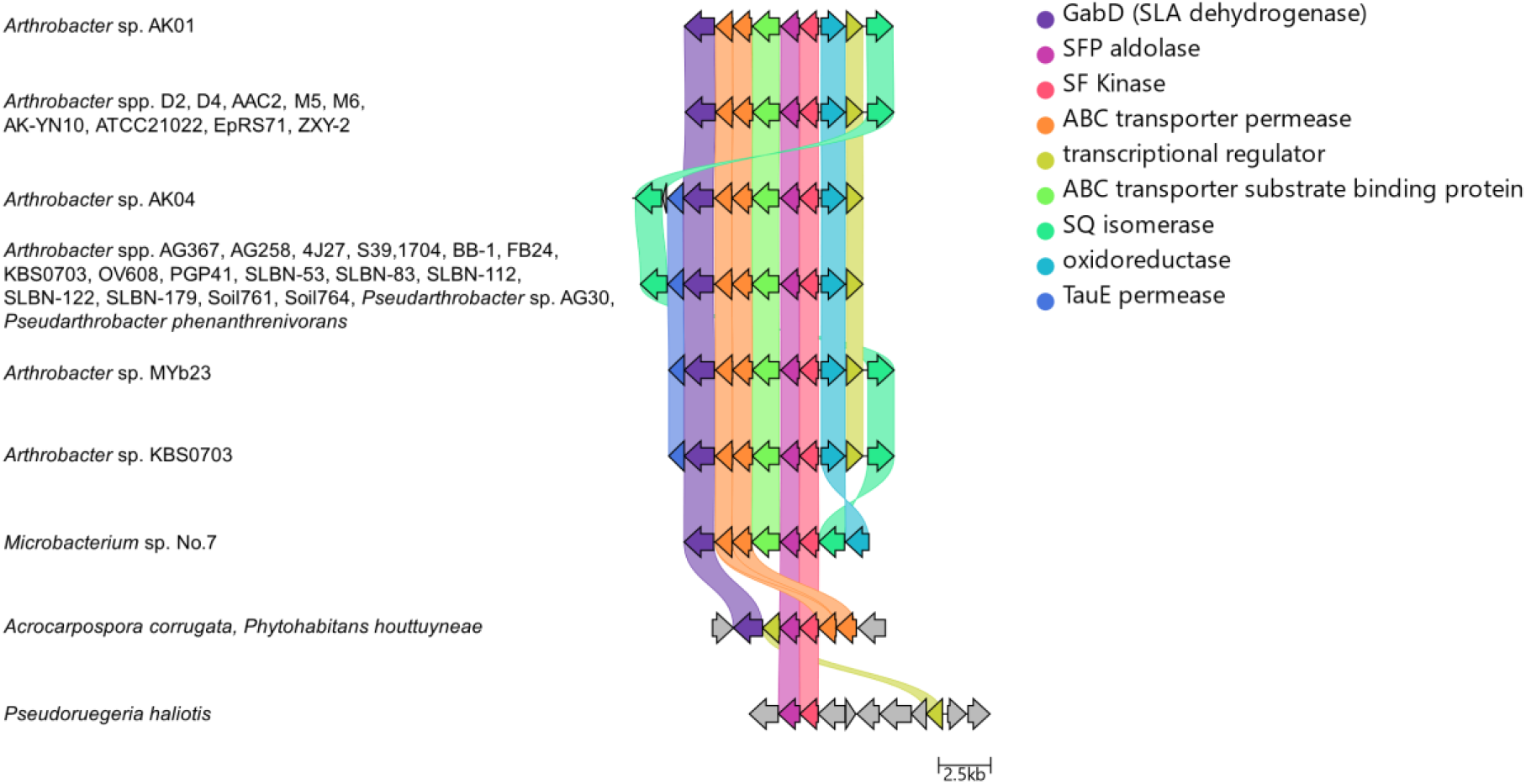
Distribution and architecture of sulfo-EMP gene clusters in *Arthrobacter* and related organisms. Syntenic relationship of sulfo-EMP gene clusters in *Arthrobacter* sp. AK01 and AK04 with homologous gene clusters. Colored links indicate ≥ 30% protein sequence similarity. Genome accession codes: *Arthrobacter* sp. D2 (LUKB01000109.1), *Arthrobacter* sp. D4 (LUKC01000078.1), *Arthrobacter* sp. AAC2 (JAAGBD010000014.1), *Arthrobacter* sp. M5 (LVCB01000107.1), *Arthrobacter* sp. M6 (LVCC01000103.1), *Arthrobacter* sp. AK-YN10 (AVPD02000157.1), *Arthrobacter* sp. ATCC 21022 (CP014196.1) *Arthrobacter* sp. EpRS71 (LNUV01000003.1), *Arthrobacter* sp. ZXY-2 (CP017421.1), *Arthrobacter* sp. AG367 (VIVE01000010.1), *Arthrobacter* sp. AG258 (SOBI01000009.1), *Arthrobacter* sp. 4J27 (CAQI01000048.1), *Arthrobacter* sp. S39 (SIHX01000007.1), *Arthrobacter* sp.1704 (SOBD01000016.1), *Arthrobacter* sp. BB-1 (VDEV01000010.1), *Arthrobacter* sp. FB24 (CP000454.1), *Arthrobacter* sp. KBS0703 (MVDG02000001.1), *Arthrobacter* sp. OV608 (FOEZ01000003.1), *Arthrobacter* sp. PGP41 (CP026514.1), *Arthrobacter* sp. SLBN-53 (VFMZ01000001.1), *Arthrobacter* sp. SLBN-83 (VFMX01000001.1), *Arthrobacter* sp. SLBN-112 (VFMU01000001.1), *Arthrobacter* sp. SLBN-122 (VFMS01000001.1), *Arthrobacter* sp. SLBN-179 (VFNR01000001.1), *Arthrobacter* sp. Soil761 (LMSF01000007.1), *Arthrobacter* sp. Soil764 (LMSI01000008.1), *Pseudarthrobacter phenanthrenivorans* (CP002379.1), *Pseudarthrobacter phenanthrenivorans* (RBNH01000003.1), *Pseudarthrobacter phenanthrenivorans* (VHJD01000009.1), *Pseudarthrobacter* sp. AG30 (QEHL01000024.1), *Arthrobacter* sp. MYb23 (PCPR01000010.1), *Arthrobacter* sp. KBS0703 (MVDG02000001.1), *Microbacterium* sp. No.7 (CP012697.1), *Acrocarpospora corrugata* (BLAD01000050.1), *Phytohabitans houttuyneae* (BLPF01000004.1), *Pseudoruegeria haliotis* (PVTD01000003.1).

## Conclusions

Two sulfoglycolytic soil bacteria belonging to the *Arthrobacter* genus (strains AK01 and AK04) were isolated by enrichment culture involving growth on SQ from soil. The stationary phase optical density of these bacteria when grown on SQ was approximately half that of growth on glucose, consistent with utilization of only three of the six carbons of SQ. Both possessed a variant of the sulfo-EMP pathway that uses an SLA dehydrogenase to produce SL that is secreted into the growth media, and which is proposed to arise from the occurrence of an SLA dehydrogenase (GabD). SL in turn becomes available for other members of the microbial community that specialize in its metabolism [23-27]. The observation that cells achieve stationary phase in SQ at approximately half the optical density as cells grown on glucose is consistent with utilization of only three of the six carbon in SQ. Prior to this work, the only sulfo-EMP pathway bacteria that have been characterized were a range of *E. coli* strains, which produce DHPS through the action of SLA reductase [5]. Notably, strains AK01 and AK04 lack genes assigned as encoding an SQase, but nonetheless could grow on MeSQ, indicating the presence of a non-specific SQase or a novel SQase that is not readily identified by sequence homology. This study highlights the core genes required for sulfoglycolysis (YihS, YihT, YihU, YihV) and constitutes the first examples of sulfo-EMP bacteria isolated from soil.

## Methods

### Bacterial growth media

Growth media was prepared using M9 minimal salts media (2 mL), trace metal solution (0.1 mL), and vitamin solution (0.01 mL) and contained 5 mM sulfoquinovose (SQ) as sole carbon source, made up to a final volume of 10 mL with water. M9 minimal salts media contains 0.45 M Na_2_HPO_4_, 0.11 M KH_2_PO_4_, 0.09 M NH_4_Cl, 0.04 M NaCl, 0.1 M MgSO_4_, 0.1 M CaCl_2_; trace metal solution contains 0.4 mM FeCl_3_, 0.08 mM CoCl_2_, 0.08 mM CuCl_2_, 0.08 mM MnCl_2_, 0.55 mM ZnCl_2_, 0.01 mM NiCl_2_, 0.05 mM Na_2_MoO_4_ and 0.5 mM H_3_BO_3_; vitamin mixture contains 0.04 mM biotin, 0.05 mM calcium pantothenate, 0.15 mM thiamine hydrochloride, 0.36 mM p-aminobenzoic acid, 0.81 mM nicotinic acid, 1.49 mM pyridoxamine dihydrochloride, 0.01 B12 (cyanocobalamin).

#### Isolation of *Arthrobacter* sp

*Arthrobacter* sp. AK01 and AK04 was isolated from enriched culture, obtained from soil of the Botany Systems Garden (University of Melbourne).

Two soil samples were collected and approximately 1 g of soil was suspended in 5 mL of sterilized growth media containing 5 mM SQ as a sole carbon source. The culture was incubated at 30 °C for 4 d with agitation at 250 rpm. A subsample (100 *μL*) was transferred into fresh vitamin supplemented M9 media and grown for a further 4 d. This step was repeated four times and after outgrowth of the final culture for 4 d, cells were plated onto LB agar plates (10 g/L tryptone, 5g/L NaCl, 5g/L yeast, 15g/L agar) and incubated overnight at 30 °C in dark. Single colonies were picked and inoculated into fresh vitamin supplemented M9 media containing 5 mM SQ and incubated at 28 °C while shaking at 250 rpm. Once the cultures were visibly turbid, cells were again plated onto LB agar, incubated overnight at 30 °C in dark and single colonies picked and inoculated again into fresh vitamin supplemented M9 media containing 5 mM SQ. Once cultures were visibly turbid, frozen stocks were prepared by diluting to 10% glycerol and freezing at -80 °C. Cell morphology was examined using scanning electron microscopy. Genomic DNA was isolated using the GenElute DNA extraction kit (Sigma) with inclusion of lysozyme and RNAase.

### Phenotypic assays

Bacteria were grown in vitamin-supplemented M9 minimal media containing ^13^C_6_-SQ (7.7 mM) at 30 °C for 1 day with agitation at 250 rpm. After cultures became visibly turbid, the cells were sedimented by centrifugation (10,000 rpm, 10 min) and the supernatant was diluted with 50 % D_2_O and transferred to a 5 mm NMR tube. ^13^C NMR spectra were acquired using a 500 MHz instrument and are shown in Figure 1(a).

### Genome sequence, assembly, and annotation

Genomic DNA was sequenced using an Illumina NextSeq at the Peter Doherty Institute for Infection and Immunity, Parkville, Victoria, Australia. DNA was prepared for sequencing on the Illumina NextSeq platform using the Nextera XT DNA preparation kit (Illumina) with ×150 bp paired end chemistry and with a targeted sequencing depth of >50×. Draft genomes were assembled using Shovill v1.1.0 (https://github.com/tseemann/shovill) and annotated using Prokka v1.14.5 [28]. GC percentage and ANI calculations were performed using ANI calculator (https://www.ezbiocloud.net/tools/ani) [29]. Assembled genomes have been deposited at the NCBI (GenBank accession: AK01, SAMN23041292; AK04, SAMN23041293). General features for isolated bacteria are reported in Table 1. The protein sequences of putative sulfo-proteins in AK01 and AK04 were used to search against the NCBI non-redundant database using BLASTp. Percentage identities for key sulfo proteins are given in **Table 4**.

### Discovery of related sulfoglycolytic operons

Sequences for *E. coli* sulfoquinovosidase (NP_418314.1, locus tag b3878), SQ mutarotase (NP_418315.3, locus tag b3879), SQ isomerase (NP_418316.4, locus tag b3880), SF kinase (NP_418319.2, locus tag b3883), SFP aldolase (NP_418317.1, locus tag b3881), SLA reductase (NP_418318.1, locus tag b3882) and sulfo-EMP regulator (NP_418320.2, locus tag b3884) were submitted separately as queries to the NCBI BLASTp tool. The database searched was the non-redundant protein sequence (nr) database, with *E. coli* (taxid: 562) sequences excluded. Standard algorithm parameters were used, except the maximum target sequences was set to 10,000. The results were filtered, with only protein sequences with E-value ≤ 5.41e-44 retained. The corresponding nucleotide accession numbers for each protein from all seven searches were extracted, and the seven lists combined and duplicates removed to give a list of candidate genome sequences. This list was converted into a MultiGeneBLAST reference library and searched using the *E. coli* sulfo-EMP gene cluster as a query. Scripts for this pipeline are available on GitHub (https://github.com/jmui-unimelb/Gene-Cluster-Search-Pipeline). Gene clusters found using this workflow were screened for clusters that contained a putative SQ isomerase, a putative SF kinase and a putative SFP aldolase, but lacked a homologous sulfoquinovosidase.

## List of Abbreviations

ABC: ATP-binding cassette
DHPS: 2,3-dihydroxypropanesulfonate
NAD(P)H: reduced nicotinamide adenine dinucleotide (phosphate)
SF: sulfofructose
SL: sulfolactate
SLA: sulfolactaldehyde
SQ: 6-deoxy-6-sulfo-D-glucose
SQase: sulfoquinovosidase
SQDG: sulfoquinovosyl diacylglycerol
SQGro: sulfoquinovosyl glycerol
Sulfo-ED: sulfoglycolytic Entner-Doudoroff
Sulfo-EMP: sulfoglycolytic Embden-Meyerhof-Parnas
Sulfo-SFT: sulfoglycolytic sulfofructose transaldolase
Sulfo-SMO: sulfoglycolytic sulfoquinovose monooxygenase
TSUP: 4-toluene sulfonate uptake permease

## Declarations

### Ethics approval and consent to participate

Not applicable

### Consent for publication

Not applicable

### Availability of data and material

The datasets generated and/or analysed during the current study are available in the GenBank database with accession codes SAMN23041292 (strain AK01) and SAMN23041293 (strain AK04). Scripts used for bioinformatic search pipeline are available on the GitHub repository, https://github.com/jmui-unimelb/Gene-Cluster-Search-Pipeline.

### Authors’ contributions

SJW conceived the study with input from SP. AK and PvdP performed enrichment culture and microbial characterization. JM synthesized 13C6-SQ. AK and JM performed bioinformatic analysis. MH and SP performed genome sequencing, assembly and analysis. AK and SJW wrote the paper with input and approval of the final draft.

