## Supplementary information for "Genome sequences of *Arthrobacter* spp. that use a modified sulfoglycolytic Embden-Meyerhof-Parnas pathway"

**Table S1. Classification and general feature of *Arthrobacter* spp. AK01 and AK04**

| MIGS ID | Property | Term | Evidence code |
|---|---|---|---|
| | Classification | Domain *Bacteria* | TAS [1] |
| | | Phylum *Actinobacteria* | TAS [2] |
| | | Class *Actinobacteria* | TAS [3] |
| | | Order *Actinomycetales* | TAS [3-6] |
| | | Family *Micrococcaceae* | TAS [3-5, 7] |
| | | Genus *Arthrobacter* | TAS [5, 8] |
| | | Species *Arthrobacter* sp. | TAS [2] |
| | | Strain: AK01, AK04 | IDA |
| | Gram Strain | Not measured | |
| | Cell Shape | Short rod-like | IDA |
| | Motility | Not reported | |
| | Sporulation | Not reported | |
| | Optimum Temperature | Not tested, used 30℃ | IDA |
| | pH range; Optimum | Not tested, used 7-8 | IDA |
| | Carbon source | Yeast extract/tryptone, glucose, sulfoquinovose | IDA |
| MIGS-6 | Habitat | Soil | IDA |
| MIGS-22 | Oxygen requirement | Aerobic | IDA |
| MIGS-15 | Biotic relationship | Free living | IDA |
| MIGS-4 | Geographic location | Melbourne, VIC, Australia | IDA |
| MIGS-5 | Sample collection | March 11, 2021 | IDA |
| MIGS-4.1 | Latitude | -37.7965449 | IDA |
| MIGS-4.2 | Longitude | 144.9595151 | IDA |
| MIGS-4.4 | Altitude | Not reported | |

*Evidence codes - IDA: Inferred from Direct Assay; TAS: Traceable Author Statement (i.e., a direct report exists in the literature)

**Table S2. Gene comparison of putative sulfo-proteins.**

| Annotation | Accession code of comparator | AK01 strain | | AK04 strain | |
|---|---|---|---|---|---|
| | | Locus tag | % Identity | Locus tag | % Identity |
| Sulfoquinovose isomerase | BAE77429.1 | AK01_00271 | 59.56 | AK04_03995 | 58.92 |
| Sulfofructose kinase | BAE77426.1 | AK01_00274 | 31.76 | AK04_03987 | 32.32 |
| Sulfofructosephosphate aldolase | BAE77428.1 | AK01_00275 | 49.66 | AK04_03988 | 50.00 |
| Succinate-semialdehyde dehydrogenase [NADP(+)] GabD | WP_016486307.1 | AK01_00279 | 49.57 | AK04_03992 | 36.74 |

* comparators BAE77429.1, BAE77426.1, BAE77428.1 are from *E. coli* and WP_01648630 is from *Pseudomonas putida* SQ1
